# The response of ETV6-NTRK3 condensates to TRK inhibitors elucidates their precise phase separation mechanism

**DOI:** 10.1101/2025.02.27.640526

**Authors:** Huan Li, Tianxin Zhu, Guangya Zhu, Jingjing Xie

**Author notes:** Correspondence (G.Z.); (J.X.). These authors contributed equally (Huan Li, Tianxin Zhu).

## Abstract

*NTRK* fusion oncoproteins are considered to mediate oncogenesis in multiple rare cancers. These fusion oncoproteins form cytoplasmic kinase condensates to hyper-activate downstream signaling pathways. However, their responses to TRK inhibitors have not been investigated. Here we show that TRK inhibitors induce a quantity increasement and liquid-to-solid transition of ETV6-NTRK3 condensates. During this process, the ETV6-NTRK3 condensates undergo rapid dissolution and subsequent reemergence, with entirely altered components. We proposed a novel phase separation model for ETV6-NTRK3 that both ETV6 and NTRK3 fragments contribute to the condensate’s formation. These findings refresh the understanding of the phase separation mechanism of NTRK fusion proteins and provide basis for phenotype-based TRK inhibitor screening strategy.

## Introduction

NTRK fusion genes resulted from chromosome translocation encodes NTRK fusion proteins^1^. NTRK fusion proteins have been identified in various tumor types^2^. Although NTRK fusion oncoproteins are relatively infrequent in common cancers^2^, they serve as the primary oncogenic drivers in certain rare cancers^**3**^. For example, the ETV6-NTRK3 fusion oncoprotein is present in more than 85% cases of breast secretory carcinoma^4^, infantile fibrosarcoma^5^, and mammary analogue secretory carcinoma (MASC)^6^. The broad oncogenic capacity of NTRK fusion oncoproteins has made their inhibitor, larotrectinib^7^, the first broad-spectrum anticancer drug to be approved solely based on its effect on a specific genetic alteration, regardless of the tumor types. Following larotrectinib, entrectinib^8^ has also been approved. Subsequent next-generation inhibitors have entered clinical trials to address resistance mutations to the first-generation inhibitors^9^.

Emerging studies have reported that abnormal protein phase separation can lead to human cancers^10-14^. Moreover, many inhibitors have been demonstrated to affect the phase separation behavior of their target proteins^12-14^. In our previous work, we identified that NTRK fusion oncoproteins undergo phase separation and form kinase condensates in the cytoplasm^15^. These condensates locally concentrate fusion proteins and their substrates, facilitating kinase reactions and sustaining the hyper-activation of downstream pathways^15^. These results indicate that phase separation is a more precise mechanism for the activation of NTRK fusion oncoproteins^15^. However, it remains unclear whether TRK inhibitors affect the phase separation behavior of NTRK fusion proteins.

In this study, we found that TRK inhibitors treatment leads to an increase in the quantity of ETV6-NTRK3 condensates, while impairing their liquid-phase properties. Combining this with the results that both the ETV6 fragment itself and the ETV6-NTRK3-K380N kinase-dead mutant form low-mobility condensates, we hypothesize that the active kinase domain is crucial for maintaining the fluidity of ETV6-NTRK3 condensates. Inhibitor treatment also results in the loss of the enrichment of downstream signaling proteins in ETV6-NTRK3 condensates, completely altering the composition of the condensates. We identified a critical time point for the transition of ETV6-NTRK3 condensates. Active condensates rapidly dissolved after TRK inhibitors treatment, and inactive condensates reassembled subsequently. Based on these experimental results, we propose a new model for the phase separation of ETV6-NTRK3. Our results provide novel insights into the activation mechanism of NTRK fusion proteins and the pharmacology of TRK inhibitors. Moreover, identifying the condensates response to inhibitors could offer new approaches for the development and optimization of inhibitors.

## Results

### TRK inhibitors treatment leads to quantity increasement and liquid-to-solid transition of ETV6-NTRK3 condensates

To investigate the response of ETV6-NTRK3 condensates to TRK inhibitors, we treated cells stably expressing ETV6-NTRK3-mEGFP with TRK inhibitors, larotrectinib and entrectinib. The results of live-cell imaging indicated that number of ETV6-NTRK3 condensates significantly increased upon treatment of both inhibitors (Fig. 1A-B). We further characterized the liquid-like properties of ETV6-NTRK3 condensates through FRAP experiments (Fig. 1C-D). Interestingly, after inhibitor treatment, the fluorescence signal of ETV6-NTRK3 phase-separated condensates showed little recovery after photobleaching (Fig. 1C-D). This result suggested that liquid-like properties of ETV6-NTRK3 condensates have been significantly impaired. These condensates underwent a transition from liquid to solid state.

**Fig. 1.**
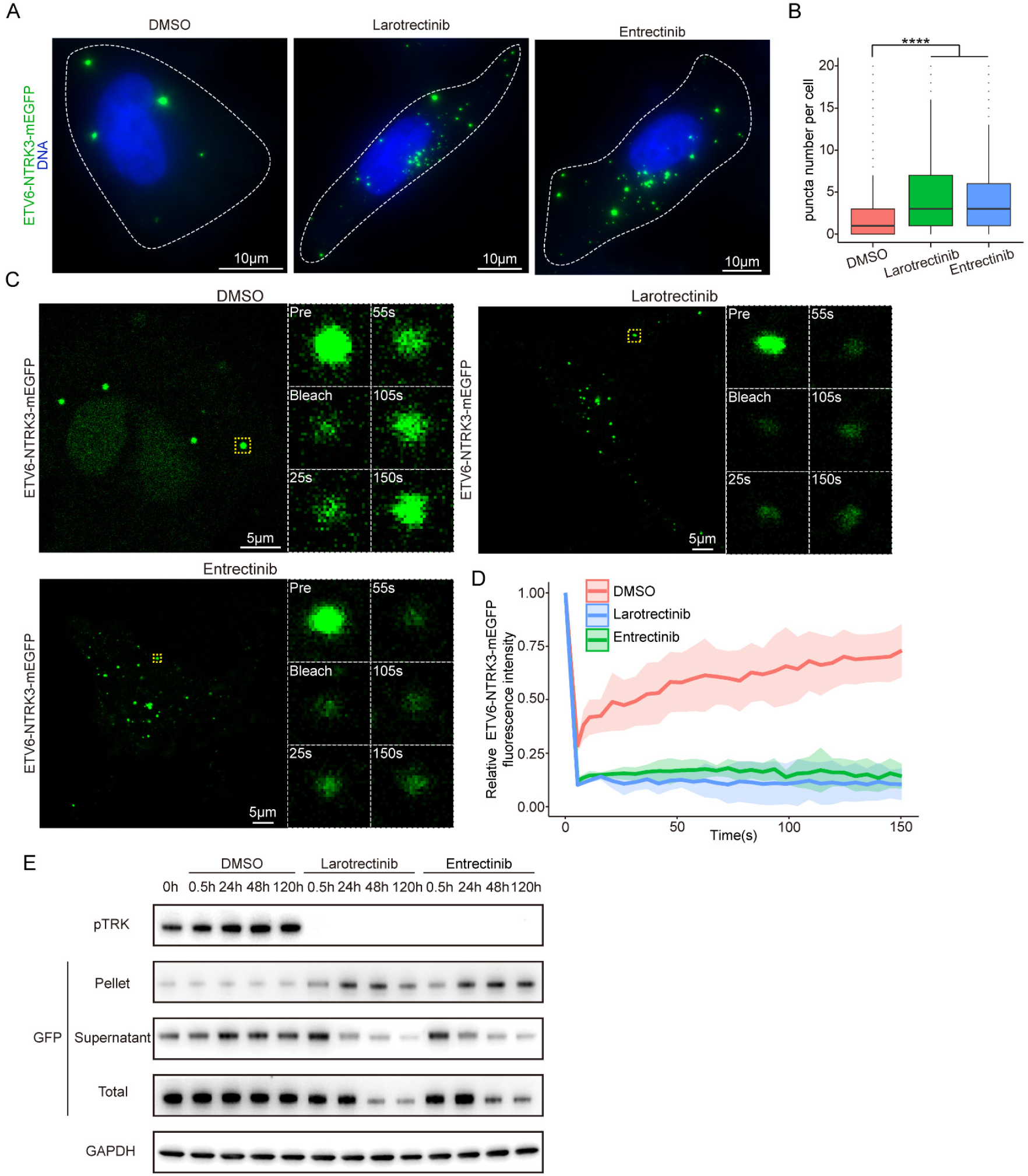
TRK inhibitors treatment leads to quantity increasement and liquid-to-solid transition of ETV6-NTRK3 condensates. (A) Representative images of ETV6-NTRK3-mEGFP merged with DAPI in stable SF268 cells treated with DMSO, Larotrectinib and Entrectinib. White dashed lines represent boundary of cells. (B) Quantification of puncta number in stable SF268 cells expressing ETV6-NTRK3-mEGFP treated with DMSO, Larotrectinib and Entrectinib. n=5000 cells. For box plots, line, median; box limits, interquartile range (IQR); whiskers, 1.5IQR. Pairwise t test, ****p<0.0001. (C) Representative series of FRAP experiments with ETV6-NTRK3-mEGFP puncta in stable SF268 cells treated with DMSO, Larotrectinib and Entrectinib. A zoomed view of FRAP series is shown on the right. (D) Quantification of FRAP experiments for ETV6-NTRK3-mEGFP puncta in stable SF268 cells treated with DMSO, Larotrectinib and Entrectinib. (means ± SD, n=3 experiments). (E) Immunoblot of the indicated proteins in stable SF268 cells expressing ETV6-NTRK3-mEGFP treated with DMSO, Larotrectinib and Entrectinib. Cells were treated with compounds for indicated duration.

Such liquid-to-solid transitions have also been observed in the phase separation processes of other proteins and are commonly regarded as an aberrant phase separation state. For example, mutations in FUS and TDP-43 proteins in amyotrophic lateral sclerosis (ALS) lead to liquid-to-solid phase transitions in their phase-separated condensates^16, 17^.

We further explored the state of ETV6-NTRK3 protein after the liquid-to-solid phase transition through immunoblots (Fig. 1E). During sample collection, soluble and insoluble components from the cells were separated by centrifugation. The results indicated that after 0.5 hour of inhibitor treatment, the phosphorylation of TRK was already suppressed (Fig. 1E), indicating that the kinase activity of ETV6-NTRK3 had been inhibited. After 24 hours of inhibitors treatment, ETV6-NTRK3-mEGFP in the soluble fraction was significantly reduced (Fig. 1E). In contrast, the content of ETV6-NTRK3-mEGFP in the insoluble fraction increased (Fig. 1E), corresponding to the liquid-to-solid phase transition of ETV6-NTRK3-mEGFP under inhibitors treatment. Additionally, the levels of ETV6-NTRK3-mEGFP in both the soluble and total fractions gradually decreased in 5 days (Fig. 1E), indicating that ETV6-NTRK3 undergoes degradation upon inhibitors treatment. This is consistent with previous reports that ETV6-NTRK3 could be degraded by larotrectinib and entrectinib^18^.

### Active kinase domains are responsible for the liquid-like properties of ETV6-NTRK3 condensates

How kinase inhibitors induce the increase in the number of ETV6-NTRK3 phase-separated condensates and their liquid-to-solid transition remained unclear. We hypothesized that the kinase domain adopts a conformation with higher interaction valency upon inhibitors binding, which enhances intermolecular interactions. The excessive interactions lead to an increase in the number of condensates and results in the sequestration of proteins inside these condensates, ultimately causing the condensates to solidify. To validate this hypothesis, we treated cells stably expressing full-length TRKC protein and the TRKC kinase domain with TRK inhibitors (SI Appendix Fig. S1A-B). The experiments were performed to observe the phase separation behavior alteration of kinase domain alone, free from the influence of ETV6 fragment. However, live-cell imaging results showed that both inhibitors failed to alter the distribution of full-length TRKC protein or the TRKC kinase domain within cells (SI Appendix Fig. S1A-B). The full-length TRKC protein maintained its membrane localization, while the TRKC kinase domain remained diffusely distributed in the cytoplasm, with no noticeable aggregation observed (SI Appendix Fig. S1A-B). This result suggested that the changes in the phase-separation state induced by kinase inhibitors may not be mediated solely through the kinase domain. The properties of the ETV6 fragment likely play an important role in this process as well.

Next, we focused on studying the ETV6 fragment itself. Based on the fusion points of ETV6 in ETV6-NTRK3, we constructed a fluorescently tagged ETV6 fragment. Live-cell imaging results showed that ETV6-mEGFP forms plaque-like aggregates in cells (Fig. 2A). These aggregates are not smooth and spherical, suggesting there may be defects in their liquid-like properties. FRAP analysis revealed that the fluorescence signal of ETV6-mEGFP aggregates shows little recovery after photobleaching, confirming their solidified nature (Fig. 2B-C). The low fluidity observed in both ETV6-mEGFP condensates and the inactive ETV6-NTRK3-mEGFP condensates suggests that the active kinase domain may be crucial for maintaining the liquid properties of ETV6-NTRK3 condensates. To verify this hypothesis, we expressed the kinase-dead ETV6-K380N-NTRK3 mutant in cells. It was reported that the kinase-activity of this mutant was completely abolished^19^. Live-cell imaging showed that ETV6-NTRK3-K380N-mEGFP formed plaque-like condensates similar to those of ETV6-mEGFP (Fig. 2D). FRAP experiments show that the mobility of ETV6-NTRK3-K380N condensates is also very low (Fig. 2E-F). These results are consistent with our expectation that the kinase domain’s activity is critical for retaining the liquid state of ETV6-NTRK3 condensates. When the kinase domain is inactive or absent, the condensates tend to transit into a solid state. This suggested that the formation of low-mobility condensates is likely attributed to the intrinsic properties of the ETV6 fragment. Previous studies have reported that the ETV6 fragments can assemble into insoluble fibers in vitro via interactions between the SAM domains^20 21^. The low-mobility plaque-like condensates of ETV6 in cells are likely a manifestation of this assembly mode. Hence, we speculated that the kinase domain may alter the assembly manner of the ETV6 fragments.

**Fig. 2.**
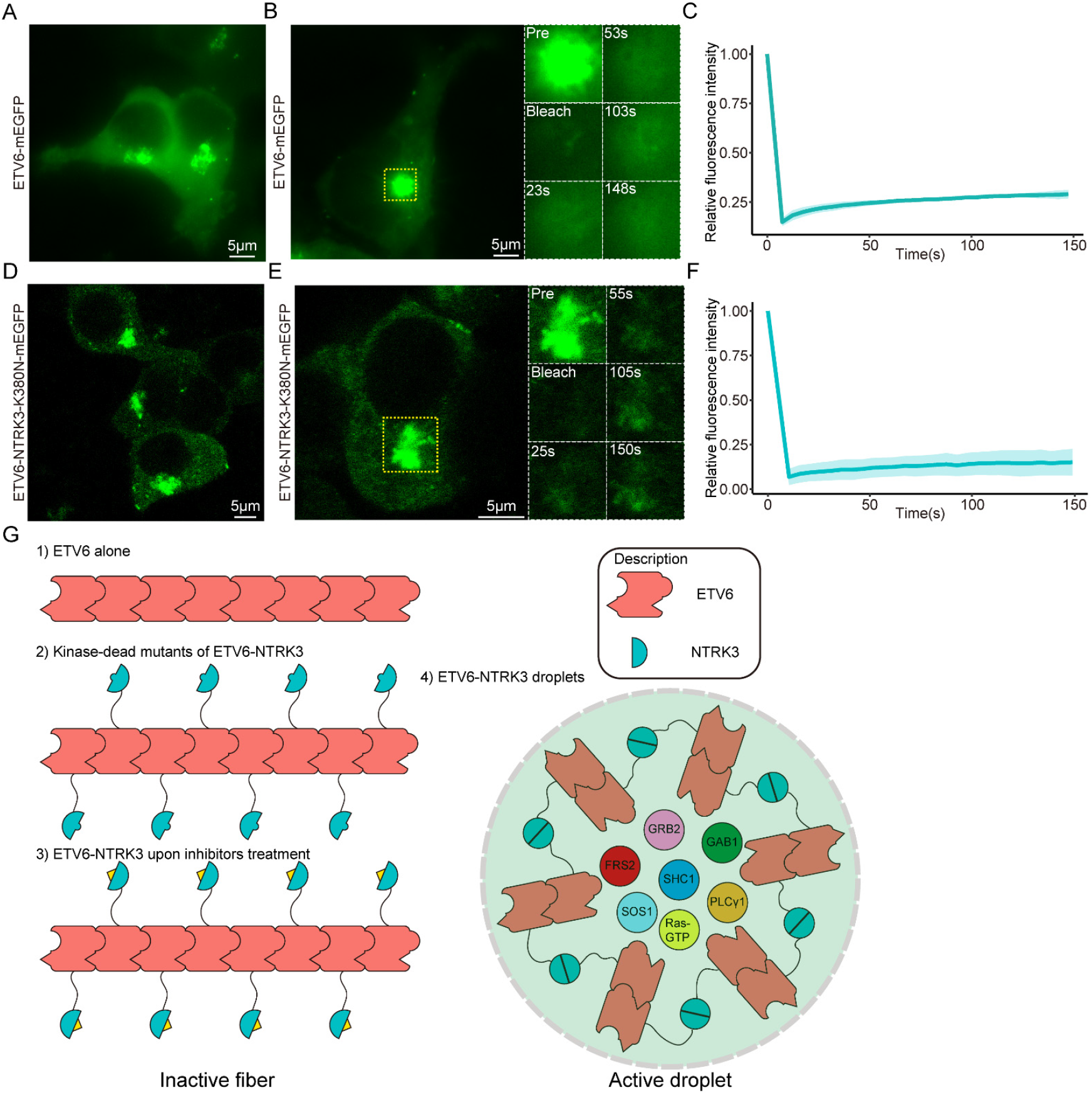
Active kinase domains are responsible for the liquid-like properties of ETV6-NTRK3 condensates. (A) Representative live cell imaging of ETV6-mEGFP in stable HEK293T cells. (B) Representative series of FRAP experiments with ETV6-mEGFP puncta in stable HEK293T cells. A zoomed view of FRAP series is shown on the right. (C) Quantification of FRAP experiments for ETV6-mEGFP puncta in stable HEK293T cells (means ± SD, n=3 experiments). (D) Representative live cell imaging of ETV6-NTRK3-K380N-mEGFP in stable HEK293T cells. (E) Representative series of FRAP experiments with ETV6-NTRK3-K380N-mEGFP puncta in stable HEK293T cells. A zoomed view of FRAP series is shown on the right. (F) Quantification of FRAP experiments for ETV6-NTRK3-K380N-mEGFP puncta in stable HEK293T cells (means ± SD, n=3 experiments). (E) Assembly model of ETV6 and ETV6-NTRK3 at different states.

Studies have shown that the transient dimerization of the active kinase domain plays a critical role in the phase separation of the EML4-ALK fusion protein^22^. This active kinase dimerization is commonly observed in various receptor tyrosine kinases^23^. Based on these findings, we proposed a model for ETV6-NTRK3 phase separation (Fig. 2G). In the absence of the active kinase domain, proteins are dominated by the ETV6 fragment and assemble into low-mobility fibers. This scenario applies to the ETV6 fragment itself, the inactive mutant ETV6-NTRK3-K380N, and ETV6-NTRK3 treated with TRK inhibitors. In contrast, the active kinase domain may disrupt fiber assembly through its own dimerization, allowing ETV6-NTRK3 to form liquid-phase condensates with downstream proteins (Fig. 2G).

### Composition of the ETV6-NTRK3 condensates has been significantly altered after TRK inhibitors treatment

Next, we characterized the internal composition of ETV6-NTRK3 condensates after inhibitor treatment. We first constructed cSH2(PLCγ1)-mEGFP to detect phosphorylation states of TRKC kinase domain. The SH2 domain were speculated to bind to phosphorylated tyrosine on the kinase domain, which only exist in the active form of kinase domain. We first tested cSH2(PLCγ1)-mEGFP in the FUS-optodroplet-NTRK3 system, and found it was rapidly recruited after the FUS-optodroplet-NTRK3 condensates formation (SI Appendix Fig. S2), demonstrating its effectiveness in detecting active TRK kinase domains. Live-cell imaging showed that cSH2(PLCγ1)-mEGFP could be recruited into the ETV6-NTRK3 condensates but was not enriched in the ETV6-NTRK3-K380N condensates (Fig. 3A). Similarly, after inhibitors treatment, cSH2(PLCγ1)-mEGFP was no longer recruited into the ETV6-NTRK3 condensates (Fig. 3A). This suggested that after inhibitors treatment, the kinase domain in the ETV6-NTRK3 condensates is no longer in its active state.

**Fig. 3.**
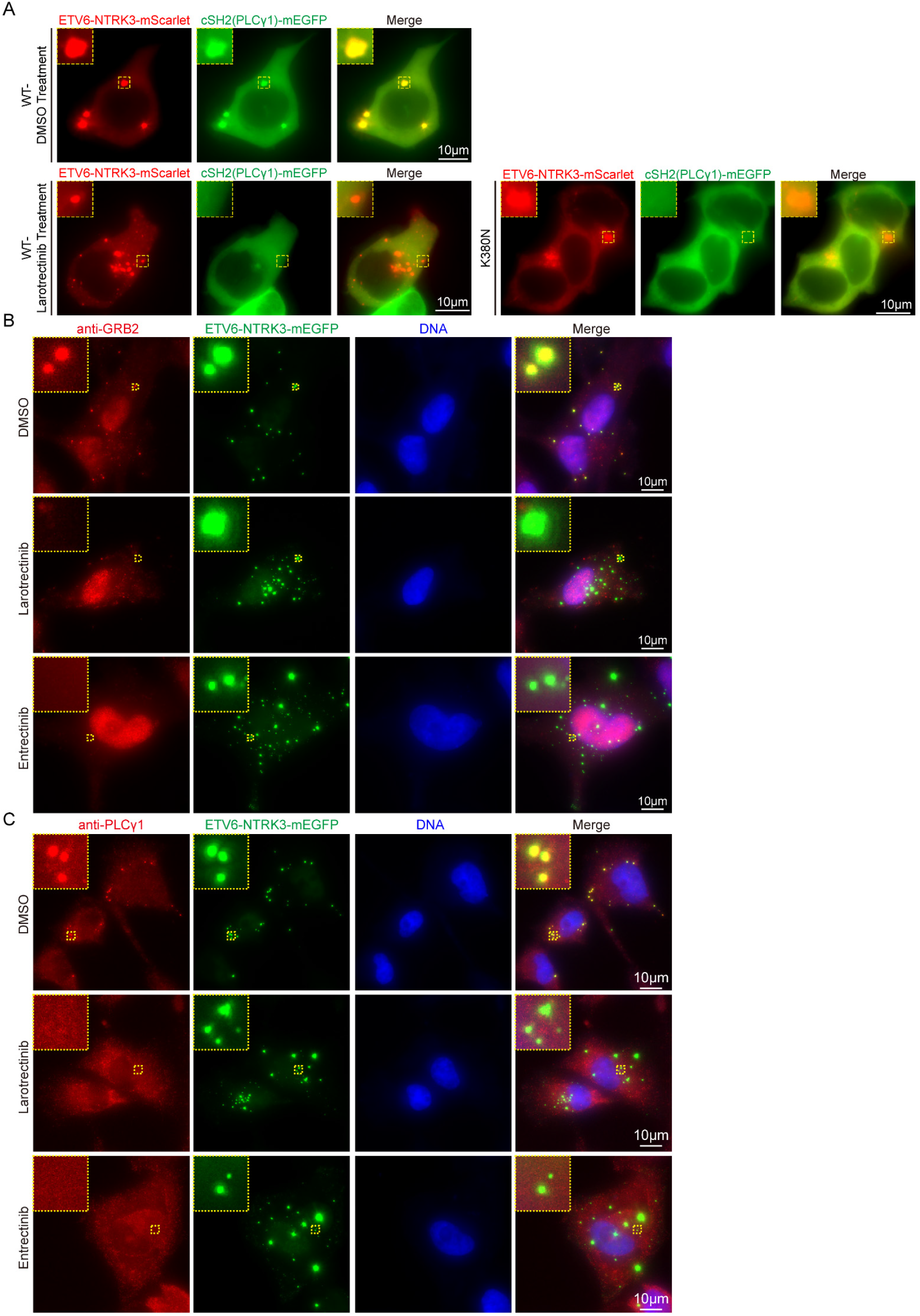
Composition of the ETV6-NTRK3 condensates has been significantly altered after TRK inhibitors treatment. (A)Representative live cell imaging of HEK239T cells expressing ETV6-NTRK3(WT/K380N)-mScarlet and cSH2(PLCγ1)-mEGFP treated with DMSO or larotrectinib. (B) Immunofluorescence imaging of GRB2 treated with DMSO, Larotrectinib and Entrectinib in stable SF268 cells expressing ETV6-NTRK3-mEGFP. (C) Immunofluorescence imaging of PLCγ 1 treated with DMSO, Larotrectinib and Entrectinib in stable SF268 cells expressing ETV6-NTRK3-mEGFP.

In previous work, we showed that downstream signaling protein GRB2 and PLCγ1 were significantly enriched in the ETV6-NTRK3 condensates^15^. However, immunofluorescence staining results indicated that GRB2 and PLCγ1 was no longer enriched in the ETV6-NTRK3 condensates after inhibitors treatment (Fig. 3B-C). This indicated that these inactive condensates have lost the ability to recruit downstream signaling proteins, and their composition has been significantly altered.

### The ETV6-NTRK3 condensates undergo rapid dissolution and subsequent reemergence after treated by TRK inhibitors

After treated with the inhibitor, the composition of ETV6-NTRK3 condensates undergoes significant changes. In combination with the observed impairment in the liquid-like properties of the condensates, it can be inferred that the condensate state has been substantially altered. However, the exact time point at which this transition occurs remains unclear. To explore this, we conducted live-cell time-lapse imaging to continuously monitor the state of the ETV6-NTRK3 condensates after inhibitor treatment. Interestingly, the addition of the inhibitor does not directly induce the increase in the number of ETV6-NTRK3 condensates. Instead, it mediated a rapid dissolution of the condensates within a short time (Fig. 4A-B). Shortly after the droplets dissolve, phase-separated condensates start to reemerge in the cytoplasm, and condensates number gradually increased (Fig. 4A-B).

**Fig. 4.**
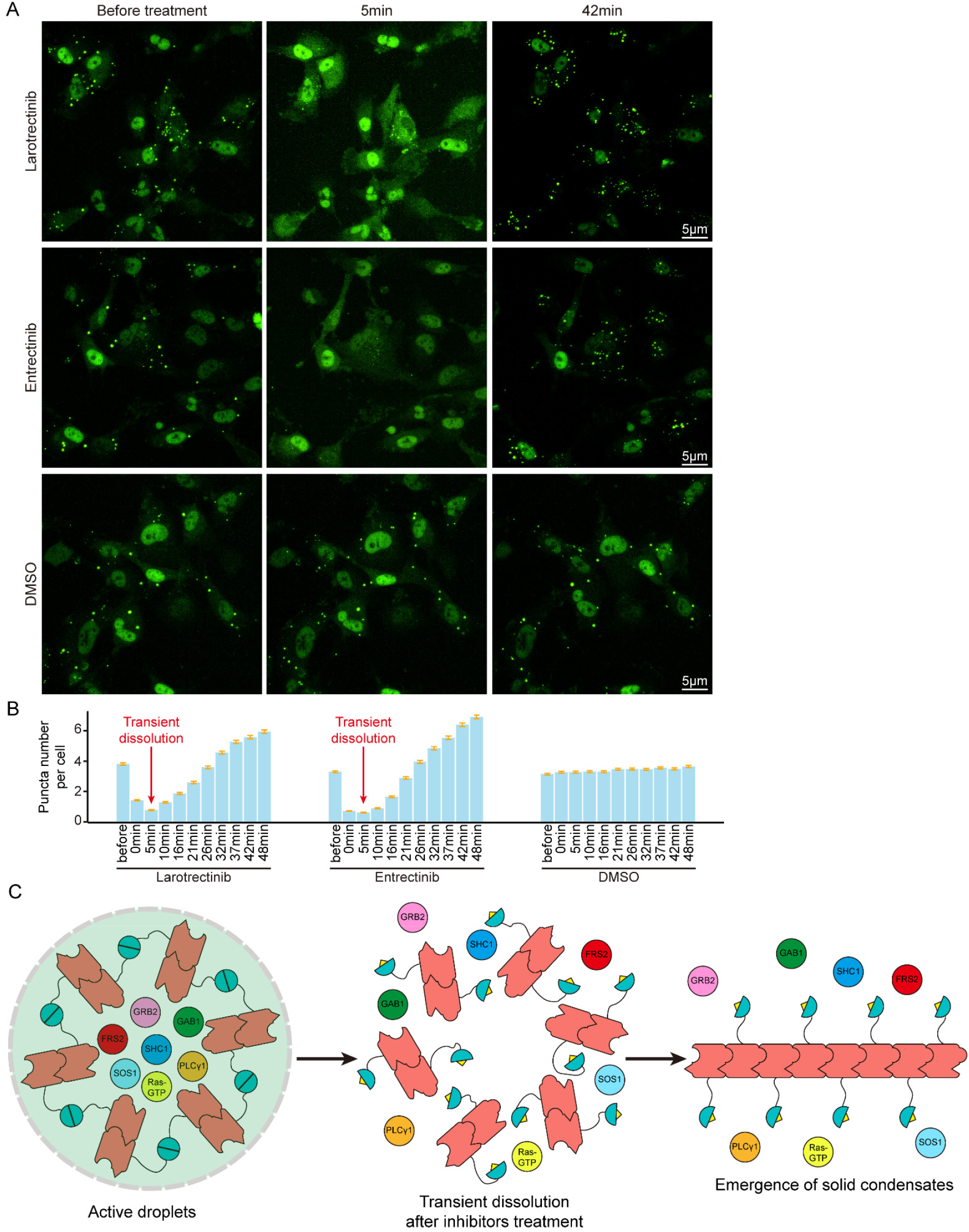
The ETV6-NTRK3 condensates undergo rapid dissolution and subsequent reemergence after treated by TRK inhibitors. (A) Representative living-cell imaging of ETV6-NTRK3-mEGFP in stable SF268 cells treated with Larotrectinib, Entrectinib and DMSO. Cells before treatment are shown on the left, cells treated for 5 minutes are shown in the middle and cells treated for 42 minutes are shown on the right. (B) Statistical analysis (means±SEM, n=3000 cells) of puncta number per cell in SF268 cells stably expressing ETV6-NTRK3-mEGFP treated with Larotrectinib, Entrectinib and DMSO. Red arrow indicates the time point when puncta number per cell reached minimum. (C) Schematic illustration of the dissolution and reemergence behavior of ETV6-NTRK3 puncta upon inhibitors treatment.

This phenomenon is consistent with the hypothesis that the assembly of ETV6-NTRK3 condensates without kinase activity follows a completely different pattern. When the inhibitor suppresses the kinase activity of ETV6-NTRK3, the original assembly mode of the condensates is disrupted, leading to the rapid disintegration of the active condensates (Fig. 4C). After this disintegration, the inactive kinase domain no longer interferes with the oligomerization of ETV6-NTRK3 Proteins begin to aggregate in a head-to-tail mode driven by the ETV6 fragment, eventually forming new ETV6-NTRK3 condensates in the cytoplasm (Fig. 4C). Due to the loss of phosphorylation in the kinase domain, the newly formed condensates no longer recruit downstream signaling proteins and exhibit overall low mobility (Fig. 4C). This model provides a comprehensive predication for the mechanism of ETV6-NTRK3 phase separation.

## Discussion

Our previous studies have thoroughly explored how NTRK fusion proteins activate themselves and downstream pathways via phase separation^15^. The phase separation of NTRK fusion proteins is mainly mediated by the N-terminal fusion partner, while the C-terminal kinase domain exhibits weak phase separation potential^15^. Therefore, TRK inhibitors targeting the kinase domain were expected to have minimal effect on the phase separation behavior of NTRK fusion proteins. However, in this study, we found that both TRK inhibitors significantly impacted the condensate state of ETV6-NTRK3, a major oncogenic NTRK fusion protein (Fig. 1A-D). This finding prompted us to further investigate the phase separation mechanism of ETV6-NTRK3 and update its phase separation model (Fig. 4C).

Aberrant protein phase separation has been identified to participate in various human cancers^10-14^. Hence, disrupting phase separation has become a novel strategy for drug development. We have reported that the SHP2 inhibitor ET070 significantly inhibit the condensates formed by SHP2 mutants^12^, and the anti-HIV drug Elvitegravir can dissolve the SRC-1 protein condensates^13^. In these cases, the inhibitors development preceded the identification of their target protein condensates. These inhibitors were initially found to inhibit the function of the target protein, and it was only subsequently discovered that they also disrupt the formation of protein condensates. These results suggest that for proteins capable of phase separation, disrupting their function is often accompanied by an impact on their phase separation behavior. Although identifying the causal relationship between protein function and phase separation is challenging, the strong correlation between them has been demonstrated. This strong correlation provides theoretical basis for phenotypic screening based on protein liquid-liquid phase separation. In 2022, Xie et al. reported an image recognition-based screening method targeting the phase separation of drug-resistant AR mutants, successfully identifying the compound ET516, which effectively inhibits drug-resistant AR mutants^14^. In 2023, Wang et al. also reported a screening method based on phase separation of FET-ETS fusion protein, identifying the positive compound LY2835219^24^. These successful cases demonstrate that the drug development strategy targeting phase-separated condensates is feasible. Here, we also demonstrated that TRK inhibitors significantly affect the phase separation behavior of ETV6-NTRK3 (Fig. 1A-D). This proves that phenotype-based screening systems based on phase separation will not miss potential kinase inhibitors. Therefore, a high-content imaging-based TRK inhibitor screening system can be established using cell lines stably expressing fluorescent protein-tagged NTRK fusion oncoproteins.

This approach not only screens for traditional kinase inhibitors but also identifies potential inhibitors that target the N-terminal fusion partner, which could provide an approach to the issue of drug-resistance mutants.

Previous studies have demonstrated that the inhibition of ETV6-NTRK3’s kinase activity leads to its degradation^18^. However, this phenomenon was not observed in TPM3-NTRK1^18^, and the mechanism remained unclear. In this study, we revealed that TRK condensates induced a liquid-to-solid transition of their condensates by inhibiting their kinase activity before inducing the degradation of ETV6-NTRK3 (Fig. 1C-E). The correlation between liquid-to-solid phase transitions and protein degradation has also been reported in cases of other proteins. For example, the estrogen receptor undergoes liquid-to-solid transition and eventual degradation upon fulvestrant treatment^25^. A similar phenomenon has been observed with PGL-1 mutants, which lead to condensate solidification and eventual degradation^26^. Cells may have specific mechanisms for clearing such aberrant solid condensates. Based on these reports, we speculate that the degradation of ETV6-NTRK3 under inhibitor treatment may also be related to the loss of condensate fluidity. Investigating this issue can offer valuable insights into the inhibitory pharmacology of TRK inhibitors against ETV6-NTRK3.

## ACKNOWLEDGMENTS

This work was supported by Shanghai Science and Technology Development Foundation (22QA1412200 to G.Z.) and the Natural Science Foundation of Shanghai (24ZR1451900 to J.X.)

## AUTHOR CONTRIBUTIONS

H.L., T.Z., G.Z. and J.X. designed the experiments; H.L., T.Z., G.Z. and J.X. wrote the manuscript; H.L. and T.Z. performed experiments; H.L. and T.Z. analyzed data.

## Methods

### Cell culture

Cells were maintained accordingly to the guidance from American Type Culture Collection (ATCC). Human HEK293T(female), HEK293FT(female) cells were cultured in DMEM supplemented with 10% (v/v) FBS, 100 units/mL penicillin and 100 mg/mL streptomycin. Human SF268(female) cells were cultured in RPMI1640 supplemented with 10%(v/v) FBS, 100 units/mL penicillin and 100 mg/mL streptomycin. HEK293T and HEK293FT cells purchased from Cobioer company have been authenticated by STR analysis and mycoplasma detection. All cells have been verified through periodic morphology checks and mycoplasma detection.

### Plasmids Transfection and viral infection

Transfection of plasmids into HEK293T/HEK293FT cells was performed using jetOPTIMUS (117-07, Polyplus-transfection SA) DNA Transfection Reagent according to the manufacturer’s instructions. Stable cells were generated via retroviral infections. Briefly, HEK293FT cells were co-transfected with viral plasmids and packaging plasmids. At forty-eight hours after transfection, the supernatant of the cell culture was collected and filtered through a 0.45 μm filter (Millipore) and used to infect cells of interest.

### Live cell imaging

Cells were grown on 24-well glass bottom plates (Cellvis, P24-1.5H-N) and images were captured with the Leica TCS SP8 confocal microscopy system using a 63× water objective (numerical aperture (NA) = 1.2) and Leica Thunder microscope using a 100× oil objective (NA = 1.44). Cells were imaged on a heated stage (37°C) and supplemented with warmed (37°C) humidified air. Fluorescent images were processed and assembled into figures using LAS X (Leica) and Fiji.

### Fluorescence Recovery After Photobleaching (FRAP)

FRAP assay was conducted using the FRAP module of the Leica SP8 confocal microscopy system and Leica Thunder microscope. The mEGFP-tagged fusion proteins were bleached using a 488-nm laser beam. Bleaching was focused on a circular or rectangular region of interest (ROI) using 100% laser power and time-lapse images were collected. Fluorescence intensity was measured using Fiji. Background intensity was subtracted and values were reported relative to pre-bleaching time points. R was used to plot and analyze the FRAP results.

### Western blot

Whole cell lysates were prepared using Cell lysis buffer for Western and IP (P0013, Beyotime). The proteins were then separated by sodium dodecyl sulfate (SDS) polyacrylamide gel electrophoresis and subsequently transferred onto a PVDF membrane (Millipore, Bedford, MA, USA). The membrane was blocked with 5% (m/v) BSA for 1 hour at room temperature and then incubated with primary antibodies at 4°C overnight followed by the incubation with secondary antibodies at room temperature. The following antibodies were used: GFP(CST#2956T), GAPDH(CST#2118S), pTRK (Tyr 674/675) (CST#4621T).

### Immunofluorescence cell staining

Cells were seeded in a 24-well glass bottom plate (Cellvis, P24-1.5H-N) and grown overnight. After 4% PFA fixation for 15 minutes at room temperature, cells were washed with PBST (1xPBS+Tween20 0.05%, 3×3min) and blocked with QuickBlock™ Blocking Buffer for Immunol Staining (Beyotime, P0260) for 30 minutes. Cells were subsequently rinsed with PBST (3×3min) and incubated with diluted GRB2 (ab32037, 1:300) or PLCγ1(CST#5690S, 1:300) antibody in QuickBlock™ Primary Antibody Dilution Buffer for Immunol Staining overnight at 4°C. After rinsing with PBST(3×3min), the cells were incubated with the diluted fluorochrome-labeled secondary antibody for 1h at room temperature followed by rinsing with PBST.

### Light activation assay of FUS-optoDroplet-NTRK3

Cells were transfected with FUS-optodroplet-NTRK3 and CSH2(PLCγ1)-mEGFP before imaging. Live cell time-lapse imaging was performed 16 hours after transfection. Light activation was performed by Leica Thunder microscopy system using a 63× water objective. A single view was imaged by alternating 488 nm and 555 nm lasers. During imaging of CSH2(PLCγ1)-mEGFP with 488nm laser, the FUS-optodroplet-NTRK3 can be activated and the optodroplets formed.

### Image data analysis

Harmony software (Perkin Elmer) was utilized to quantitatively evaluate puncta number of NTRK fusion oncoproteins and their mutants. 63× and 20× water immersion lens (Operetta CLS, Perkin Elmer) was used to acquire images. All images were captured in the confocal mode. For puncta analysis, we combined built-in building blocks to find nucleus, segment cells, find spots and do measurements. Find Spot module was used to identify all potential spots including low-contrast spots. Then a self-trained classifier utilizing built-in PhenoLOGIC module through manually labeling some true spots and false spots was applied to filter out negative spots. The spots used for classifier training were covered with as much diversity of spot intensity, size and shape as possible. The measured properties of cells and spots were finally exported. Further statistics and plotting were performed in R.

### Statistical analysis

Statistical analyses were done using R. Differences of means were tested for statistical significance with one-way analysis of variance (ANOVA) for three and more groups comparisons. For ANOVA analyses, Tukey HSD was used for multiple comparisons correction. For For box plots: line indicates median; box limits indicate interquartile range (IQR); whiskers indicate 1.5 IQR; and points indicate outliers (3 IQR). A P<0.05 was considered statistically significant.

### Materials availability

Plasmids, compounds, and cell lines generated in this study will be made available upon request. We may require a payment and/or a completed Materials Transfer Agreement in case there is potential for commercial application.

## Data availability

This manuscript includes all data involved this study. No unique code was generated in this study.

