## Supplementary Information Appendix for "The response of ETV6-NTRK3 condensates to TRK inhibitors elucidates their precise phase separation mechanism"

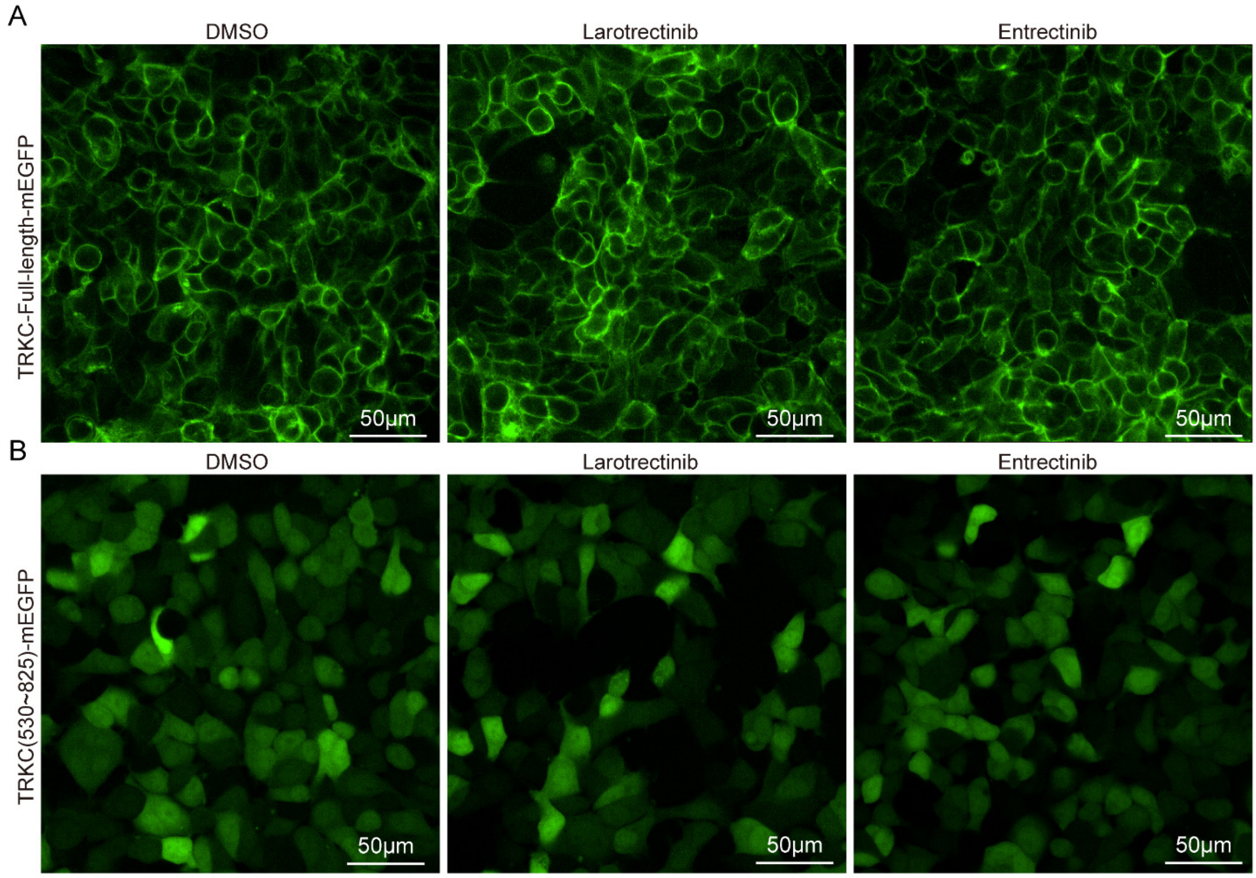


**Figure S1. TRK inhibitors generated little effect on full length TRK proteins and TRK kinase domains**

(A) Representative live cell imaging of NTRK3-Full-length-mEGFP in stable HEK293T cells treated with DMSO, Larotrectinib and Entrectinib. Cells were treated with compounds for 2 hours.

(B) Representative live cell imaging of NTRK3(530~825)-mEGFP in stable HEK293T cells treated with DMSO, Larotrectinib and Entrectinib. Cells were treated with compounds for 2 hours.

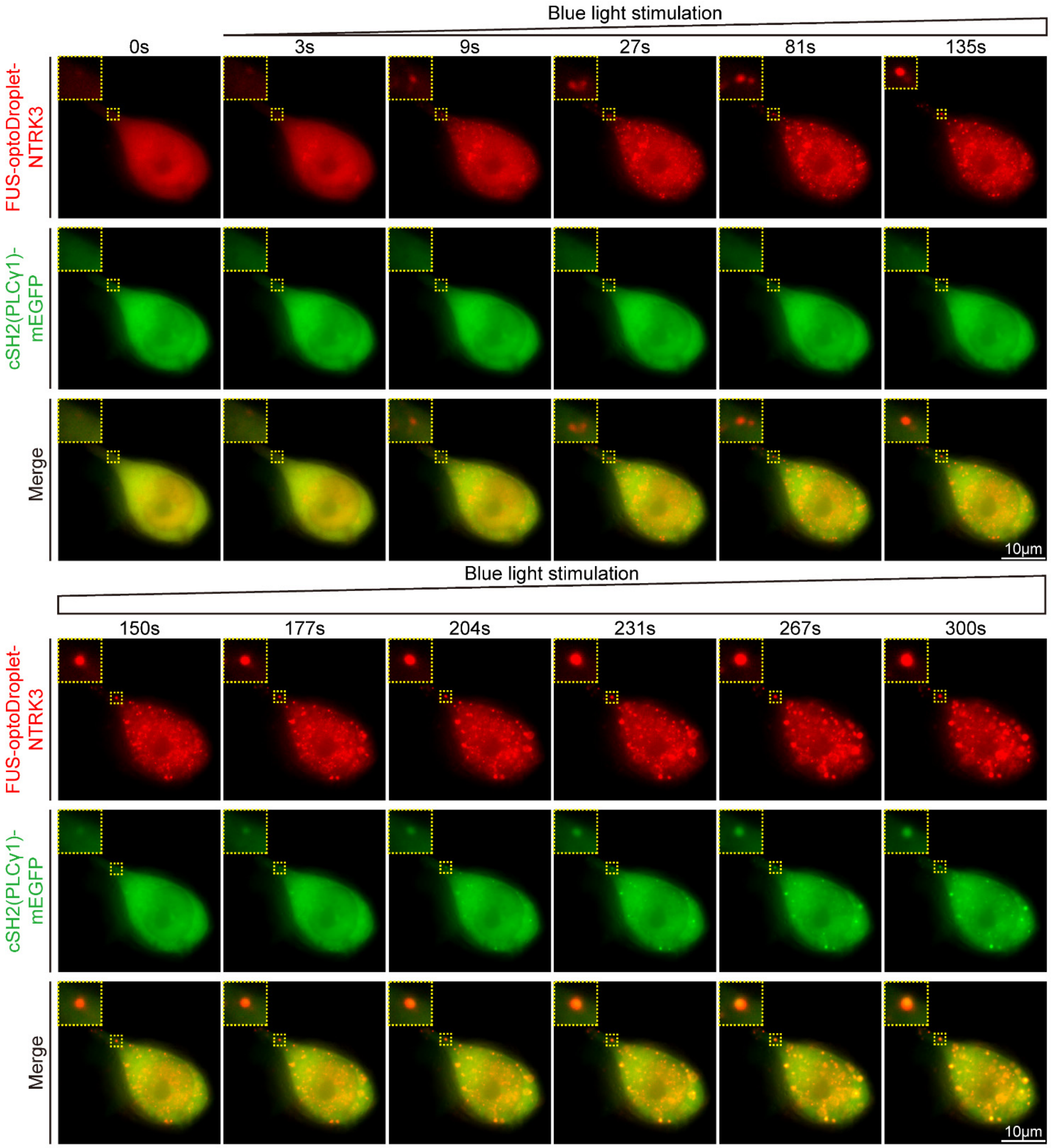


**Figure S2. CSH2(PLCγ1) probed kinase activation in FUS-optodroplet-NTRK3 condensates**

Live cell time-lapse imaging of HEK293T cells co-expressing FUS-optodroplet-NTRK3 and cSH2(PLCγ1)-mEGFP.
